# Cerebellum models of psychosis implicate association nuclei in the pathogenesis of psychosis and mechanisms of cognitive impairment

**DOI:** 10.1101/2020.11.06.372300

**Authors:** Xuebin Chang, Xiaoyan Jia, Debo Dong, Yulin Wang

## Abstract

To comprehensively investigate the white matter (WM) features of cerebellum in patients with schizophrenia, and further assess the correlation between altered WM features and clinical and cognitive assessments. Forty-two patients and fifty-two matched healthy controls (HCs) of the Collaborative Informatics and Neuroimaging Suite Data Exchange tool were involved in this study. The cerebellar WM volume was calculated by voxel-based morphometry. And tract-based spatial statistics was used to analysis the diffusion changes in patients when compared to HCs. Furthermore, we investigated the correlation between altered imaging feature and clinical, cognitive assessments. Compared to HCs, the schizophrenia patients did not reveal difference in cerebellar WM volume and schizophrenia patients showed decreased fractional anisotropy and increased radial diffusivity in left middle cerebellar peduncles and inferior cerebellar peduncles in voxel-wise but not in tract-wise. Critically, these cerebellar changes were associated with disease duration in schizophrenia patients. And significant correlation between the altered cerebellar WM features and cognitive assessments only revealed in HCs but disrupted in schizophrenia patients. The present findings suggested that the voxel-wise WM integrity analysis might was the more sensitive way to investigate the structural abnormalities in schizophrenia patients. Middle cerebellar peduncles and inferior cerebellar peduncles may be a crucial neurobiological substrate of cognition and thus might be regarded as a biomarker for treatment.

## Introduction

Our understanding of the contribution of the cerebellum to neural function and structure has changed from a traditional view centered on motor coordination to a modern understanding, and the cerebellum has also been involved in a wide range of high-level cognitive and emotional processes [1, 2]. Increasing evidence also supports the involvement of the cerebellum in multiple mental illnesses such as schizophrenia (SZ) [1, 3–5]. To date, most psychiatric studies that study the role of the cerebellum have been conducted based on classification diagnostic criteria, which treat mental illness as an independent entity [6]. It is increasingly recognized that the existing clinical diagnostic categories may not be the best because of the large overlap of symptoms, cognitive dysfunction, and genetic factors of multiple psychosis [7]. These overlaps can be reflected by shared neurobiological structures and functions and polymorphic abnormalities in psychiatric syndromes [7, 8].

Recent clinical neuroscience research has begun to use referral methods to emphasize the importance of cerebellar microstructural changes in a wide range of psychopathological risks [9, 10]. Previous animal and human neuroimaging studies have provided converging evidence for the involvement of cerebellar function in a wide range of behaviors that are dependent on with multiple cerebral cortical regions [4, 10, 11]. Accumulating evidence supports, after the discovery of focal lesions of the cerebellum, the patient showed a series of cognitive deficits, including impaired executive function, spatial cognition, language processing, emotional regulation and personality changes[12]. Since then, advanced cognitive dysfunction related to abnormal structure and function of the cerebellum has been found in many neurological diseases [13, 14].

Studies on white matter changes in SZ have focus primarily on cerebral white matter tracts [15], although the white matter of the cerebellum seems to change [16]. The extent and location of cerebellar white matter changes in SZ, and whether these changes are different in SZ variants, but it is unclear. The output fibers of the cerebellum (excluding the vestibular cerebellum to the vestibular nucleus) mainly originate from the four deep cerebellar nuclei: the dentate nucleus, the embolic nucleus, the globular nucleus and the parietal nucleus. In addition, the inferior cerebellar peduncles (ICP) contains efferent connections from the cerebellum to the vestibular nuclei [17, 18].All input fibers of the cerebellum need to pass through the middle cerebellar peduncles (MCP) [17]. However, to date, there has been no comprehensive study on the structural integrity of the SZ cerebellar white matter tract or its relationship with cognitive function.

So far, diffusion tensor (DTI) research in neuropsychiatric diseases has almost completely focused on the long-distance areas within the white matter. In patients with schizophrenia, these studies mainly found changes in the frontotemporal, interhemispheric, and frontal thalamic white matter tracts [19, 20], inferred mainly from reductions in fractional anisotropy (FA), a measure of directional dependence of diffusion of water, and exaltations in mean diffusivity (MD), axial diffusivity (AD) and radial diffusivity (RD). These results have been generally interpreted as supporting the overall theory of schizophrenia, which is considered to be a disconnection syndrome, because the white matter area forms a matrix of remote cortical white matter connectivity [21]. Decreased FA in brain white matter area may cause cognitive dysfunction in patients with schizophrenia[22, 23]. However, little is known about cerebellar white matter diffusion indicators in patients with schizophrenia.

The purpose of this study is to determine the pattern and severity of cerebellar white matter changes in SZ group, and to determine its association with cognition. We supposed that both the cerebellar infarcts and cerebellar peduncles (i.e., ICP and MCP) connecting the cerebellum and cerebellum will be affected and related to common cognitive features in SZ.

## Materials and methods

### Participants

This study included 42 schizophrenia patients and 52 healthy controls (HCs). The imaging and phenotypic information of data was downloaded from the Collaborative Informatics and Neuroimaging Suite Data Exchange tool (COINS; http://coins.mrn.org/dx) [24] and data collection was performed at the Mind Research Network, funded by a Center of Biomedical Research Excellence (COBRE) grant from the National Institutes of Health. The diagnostic confirmation of schizophrenia was confirmed by the Structured Clinical Interview for DSM-IV Axis I Disorders. Psychopathological symptoms of schizophrenia were evaluated using the Positive and Negative Syndrome Scale (PANSS) [25]. All patients were treated with antipsychotics and the antipsychotic medication was converted to chlorpromazine equivalents. The MATRICS Consensus Cognitive Battery (MCCB) cognitive battery of all participants was also included in this study. All participants were excluded for a history of substance abuse or dependence within the last 12 months, a history of neurological illness, and traumatic brain injury. Written informed consent was obtained from all participants according to institutional guidelines required by the Institutional Review Board at the University New Mexico (UNM). Five patients and three HCs were excluded for the T1 and/or scanning was not included all cerebellum. Finally, 37 schizophrenia patients and 49 HCs were included in the final analysis. The detailed demographic, clinical, and cognitive information of all patients and HCs are shown in Table 1.

**Table 1:** Demographic characteristics of the schizophrenia patients and healthy controls

| Variables | Patients (n = 37) |  | Healthy controls (n = 49) |  | P value |
| --- | --- | --- | --- | --- | --- |
|  | Mean | SD | Mean | SD |  |
| Age (years) | 38.73 | 13.79 | 38.90 | 12.07 | 0.952 |
| Gender (male: female) | 28: 9 |  | 36: 13 |  | 0.816 |
| Handedness (right: left: both) | 34: 2: 1 |  | 45: 2: 2 |  | 0.907 |
| Processing Speed | 34.51 | 11.59 | 53.73 | 8.14 | <0.001 |
| Attention Vigilance | 33.86 | 13.73 | 50.36 | 9.91 | <0.001 |
| Verbal Working Memory | 37.46 | 13.70 | 48.22 | 11.08 | <0.001 |
| Verbal Learning | 37.86 | 8.36 | 45.02 | 6.59 | <0.001 |
| Visual Learning | 35.43 | 11.64 | 46.84 | 9.85 | <0.001 |
| Reasoning Problem Solving | 42.00 | 10.25 | 54.70 | 7.66 | <0.001 |
| Social Cognition | 40.35 | 11.97 | 52.78 | 9.78 | <0.001 |
| Overall Composite | 29.25 | 12.83 | 49.74 | 8.98 | <0.001 |
| Chlorpromazine equivalents (mg/d) | 396.78 | 354.14 | - | - |  |
| Duration of illness (years) | 18.19 | 13.77 | - | - |  |
| PANSS-positive | 14.35 | 4.60 | - | - |  |
| PANSS-negative | 15.03 | 5.45 | - | - |  |
| PANSS-general | 29.35 | 8.07 | - | - |  |
| PANSS-total | 58.73 | 13.71 | - | - |  |

### Data acquisition

All images were collected on a 3-T Siemens Trio scanner with a 12-channel radio-frequency coil at the Mind Research Network. High resolution T1-weighted structural images were obtained using a five-echo MPRAGE sequence with following imaging parameters: time of repetition (TR) = 2.53 s, echo time (TE) = 1.64, 3.5, 5.36, 7.22, 9.08 ms, inversion time (TI) = 1.2 s, flip angle = 7°, filed of view (FOV) = 256 × 256 mm, number of excitations = 1, slice thickness = 1mm. The scan parameters of DTI were as follows: TR = 9 s; TE = 84 ms ; field of view (FOV) = 256 × 256 mm; slice thickness = 2 mm; number of slices = 72; slice gap = 2 mm; voxel resolution 2 × 2 × 2 mm; flip angle =90° ; number of diffusion gradient directions = 35, b = 800 s/mm^2^. All images were registered to the first b = 0 image.

### VBM analysis

To investigate the structural morphological characteristics of cerebellar white matter (WM) in patients with schizophrenia, the cerebellar-specific voxel-based morphometry (VBM) analysis was performed using the Spatially Unbiased Infratentorial template (SUIT, http://www.diedrichsenlab.org/imaging/suit.htm) toolbox implemented in Statistical Parametric Mapping, Version 12 (SPM 12, https://www.fil.ion.ucl.ac.uk/spm/software/spm12/). Before the calculation of VBM, quality control of T1 images was carried out and subjects without a complete cerebellar scan were excluded in the subsequent analysis. The steps of VBM analysis were as following. First, individual T1-weighted sequences were manually reoriented the image origin at the anterior commissure. Next, the segment and isolate function of SUIT was used to isolate the infratentorial structure (cerebellum and stem) from the surrounding tissue and segment the infratentorial structure into WM, grey matter, and cerebrospinal fluid. Then, the individual WM was normalized to the SUIT space using the Diffeomorphic Anatomical Registration Through Exponentiated Lie Algebra (DARTEL) algorithm and modulated by the deformation fields to preserve the original volume of the tissue. Finally, the resulted WM volume maps were smoothed using a 6 mm full width at half-maximum (FWHM).

### DTI analysis

To investigate the structural diffusion features of cerebellar WM in patients with schizophrenia, the analysis of DTI data was carried out using the FMRIB Software Library (FSL, www.fmrib.ox.ac.uk/fsl). First, nonbrain tissues were removed from the DTI data using the brain extraction tool (BET) algorithm in FSL. Next, head motion and eddy current corrections were carried out by the affine transformation between the gradient images and the baseline b = 0 image. Then, diffusion tensors were calculated using dtifit tool in FSL, and subsequently, FA, MD, AD, and RD maps were obtained. Besides, all subjects’ FA maps were aligned with the Montreal Neuroimaging Institute (MNI 152) template space by using the non-linear registration tool FNIRT. Furthermore, the deformation fields from FA maps were used to project the registered MD, AD, and RD maps onto the FA skeleton. Finally, the resulted maps were smoothed using a 6 mm FWHM.

### Statistical analysis

The group difference about VBM between patients and HCs was calculated by permutation-based non-parametric testing with 5000 permutations which was applied using the threshold-free cluster enhancement method, and brain volume was regressed out as covariate.

In terms of statistical analysis about voxel-wise DTI features, a voxel-wise general linear model (GLM) was used for between group comparisons of cerebellar FA map integrity. Permutation-based non-parametric testing with 5000 permutations was applied using the threshold-free cluster enhancement (TFCE) method [26]. The statistical maps of the analyses were binarized at the threshold of p < .05, family wise error (FWE) corrected for multiple comparisons. Then, the binarized maps were multiplied to create cerebellar WM masks to determine WM changes within the cerebellum. In addition, the similar processing and statistics also carried out in MD, AD, and RD map. Results with a cluster extent threshold of 100 contiguous voxels are reported.

In terms of statistical analysis about tract-wise DTI features, we used the probabilistic atlas of cerebellar WM in the MNI152 space and created masks of three pairs of cerebellar peduncles [27]. The FA map were then multiplied to create inclusive masks with the masks of cerebellar peduncles to identify the location and severity of changes in the three cerebellar WM tracts [28]. The average FA values from each tract was extracted by averaging all voxels belonging to the tract. The FA values of each tract were analyzed using the Student’s t-test with group (SZ subjects and HCs) [29] In addition, the similar processing and statistics also carried out in MD, AD, and RD map.

Finally, to investigate the correlation between altered WM features of cerebellum and the cognition assessments in both patient group and HCs group, we calculated the Spearman correlations between the cognition assessments and WM features. Besides, we also performed the Spearman correlation between the clinical indicators and altered WM features, which demonstrated significant alterations in the patient group and clinical indicators.

## Results

### VBM analysis

To investigate the structural morphological differences of cerebellar WM between schizophrenia patients and HCs, we contrasted the cerebellar WM volume maps between two groups. The schizophrenia patients did not differ from HCs regarding the cerebellar WM volume.

### DTI analysis

In voxel-wise DTI features, compared to HCs, schizophrenia patients showed widespread WM changes in left MCP and ICP. In detail, schizophrenia patients showed decreased FA (MNI coordinate: −12, −40, −41) and increased RD in left MCP and ICP (MNI coordinate: −14, −43, −38) (p < 0.05, FDR corrected) (Figure 1). The schizophrenia patients did not differ from HCs regarding MD and AD.

**Figure 1.**
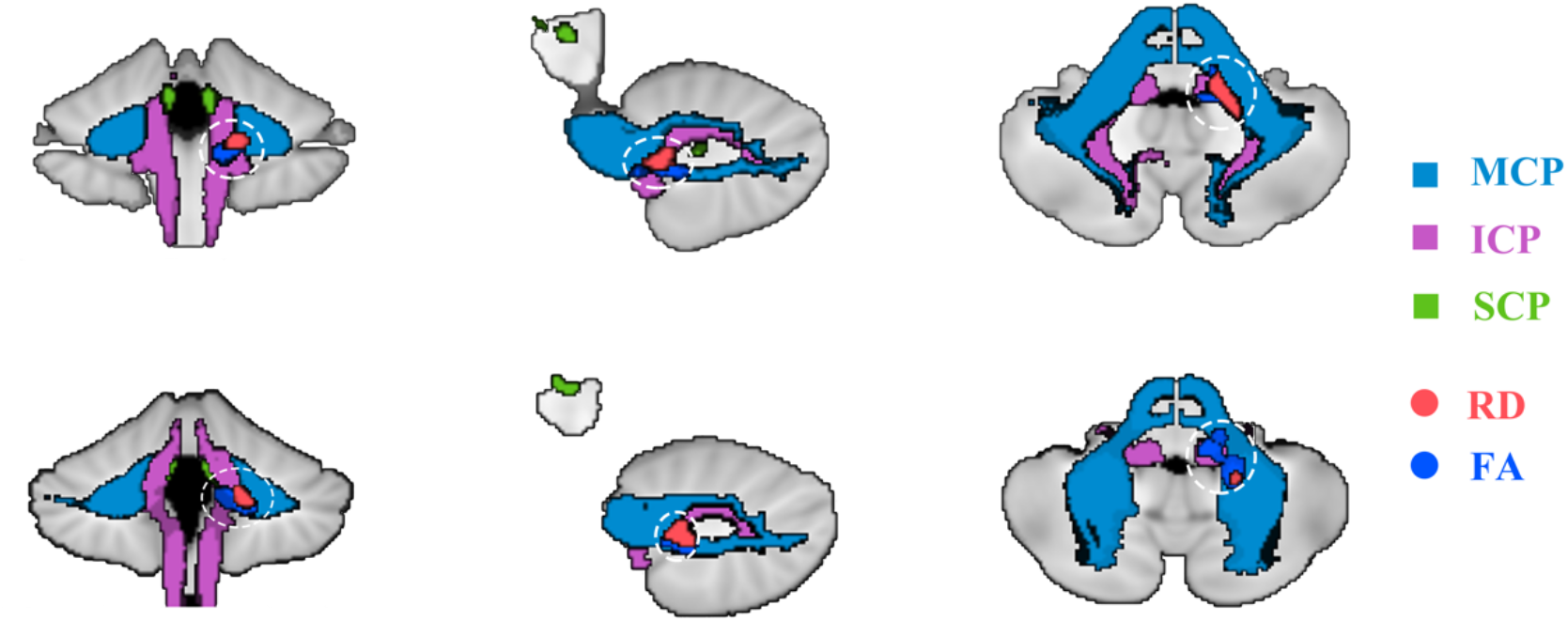
Significant group difference about FA and RD between patients and healthy controls.

In tract-wise DTI features, no significant differences were found between schizophrenia patients and HCs in any DTI features.

### Correlations between altered WM features and clinical and cognitive assessments

For the correlations between altered WM features and clinical assessments, the mean FA value in the altered region was negatively correlated with disease duration (r = −0.453, p = 0.005, Figure 2A), and the mean RD value in the altered region was positively correlated with disease duration (r = 0.647, p < 0.001, Figure 2B).

**Figure 2.**
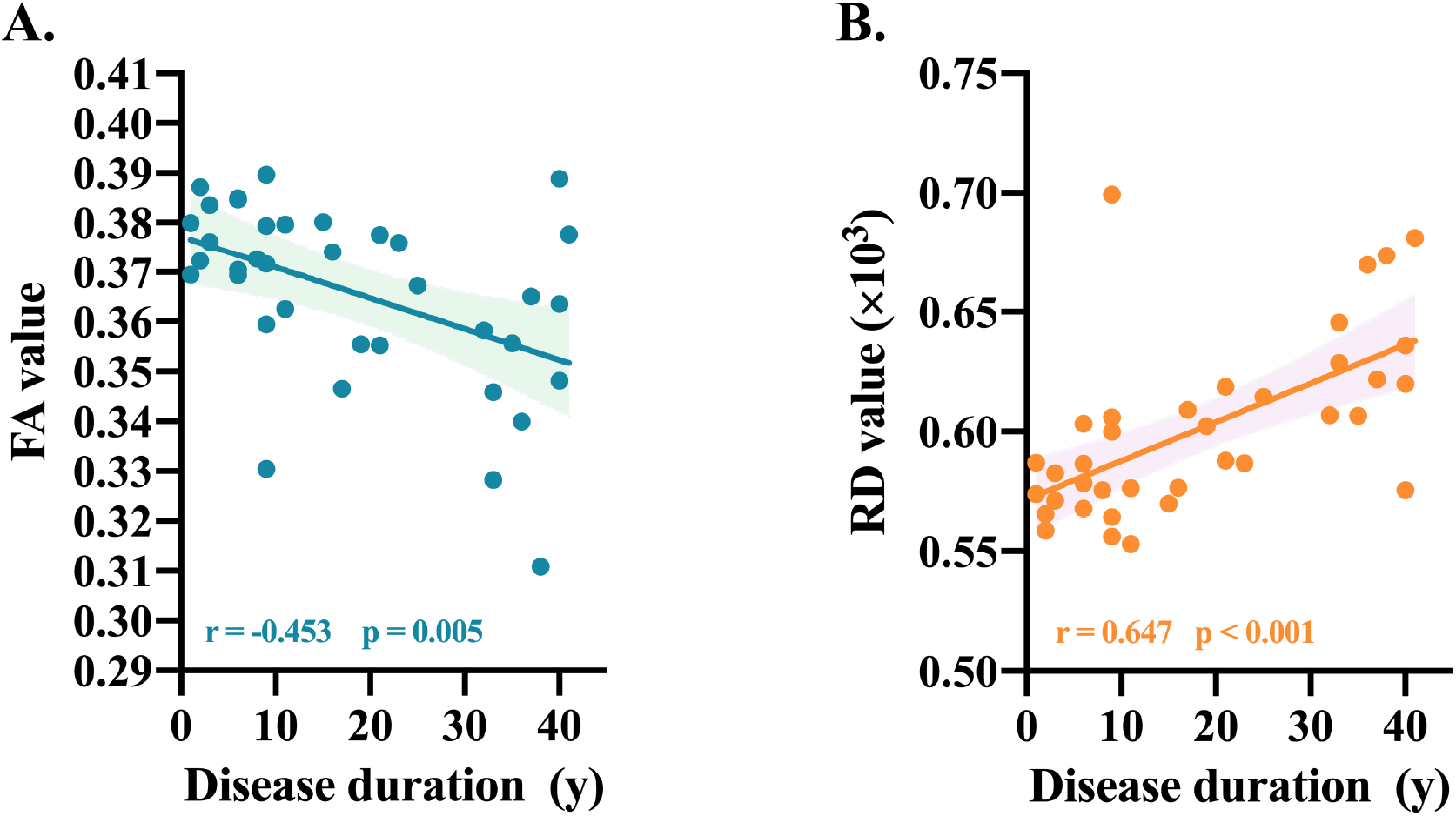
The correlation between altered diffusion features and disease duration.

For the correlations between altered WM features and cognitive assessments, significant positive correlation was showed between the mean FA value in the altered region and cognitive assessments in HCs but not showed in schizophrenia patients. Similarly, significant negative correlation was showed between the mean RD value in the altered region and cognitive assessments in HCs but not showed in schizophrenia patients. In detail, the mean FA value in the altered region in HCs was positively correlated with processing speed (r = 0.486, p < 0.001, Figure 3A), working memory (r = 0.370, p = 0.009, Figure 3B), overall composite (r = 0.370, p = 0.015, Figure 3C), and attention vigilance (r = 0.334, p = 0.025, Figure 3D), but no significant correlation was showed in schizophrenia patients (Figure 3). Besides, the mean RD value in the altered region in HCs was negatively correlated with attention vigilance (r = −0.296, p = 0.048, Figure 3E), but no significant correlation was showed in schizophrenia patients (Figure 3).

**Figure 3.**
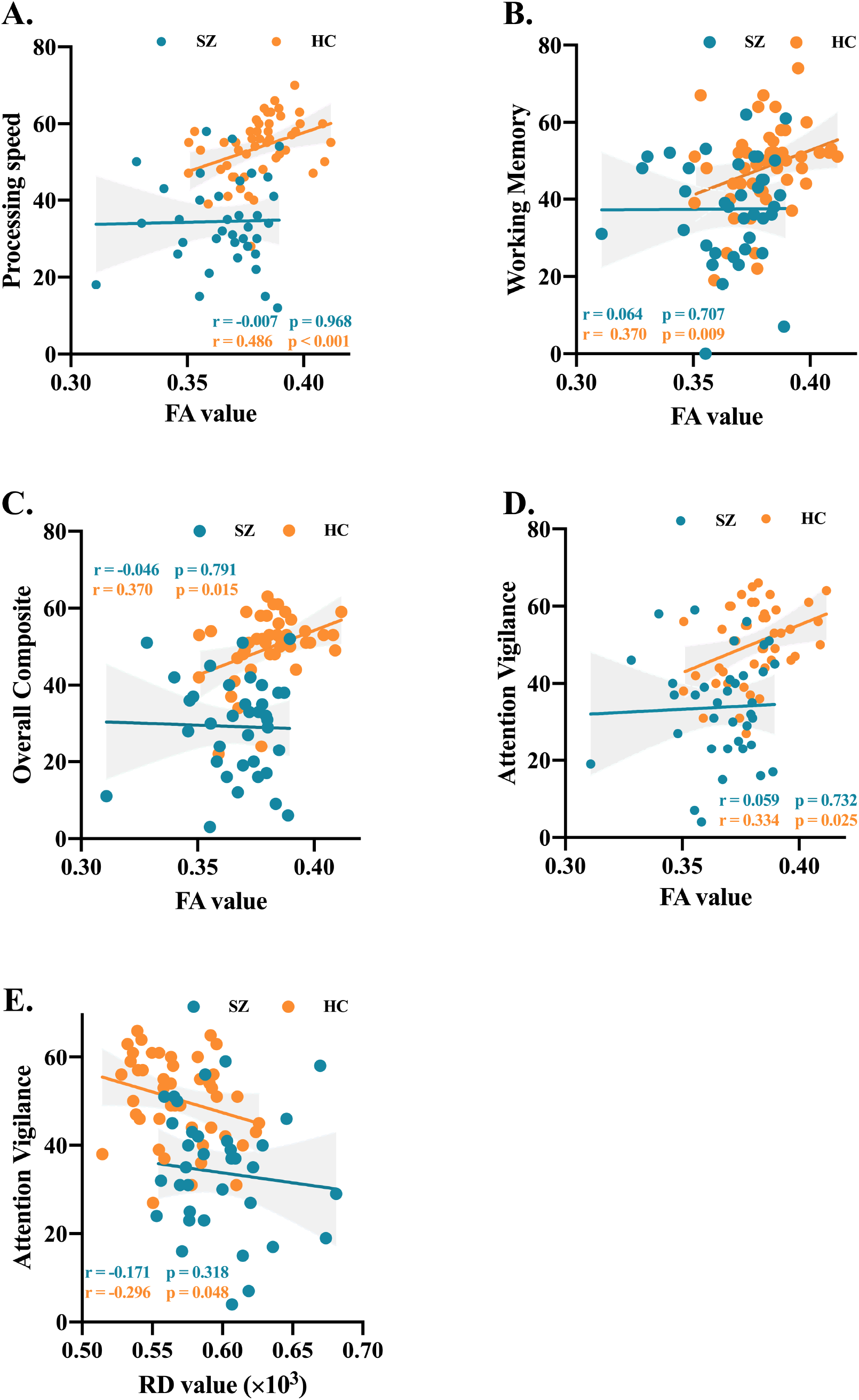
The correlation between altered diffusion features and cognitive assessments.

## Discussion

To the best of our knowledge, this is the first study to comprehensively investigate the WM features of cerebellum in patients with schizophrenia, and further assess the correlation between altered WM features and clinical and cognitive assessments. The key findings of this study were widespread voxel-wise WM abnormalities (FA and RD) involving the left MCP and ICP. Critically, these cerebellar changes were associated with disease duration in schizophrenia patients and significant correlation between the altered WM features and cognitive assessments only revealed in HCs but not in schizophrenia patients. The present findings suggested that the voxel-wise WM integrity analysis might was the more sensitive way to investigate the structural abnormalities in schizophrenia patients. Interestingly, the altered WM integrity was associated the disease duration and related to the cognitive dysfunction in schizophrenia patients.

Previous studies have investigated the WM structural connectivity [30–32] or voxel-based morphometry [33–35] in whole brain in schizophrenia patients. However, no study has comprehensively focused on cerebellar WM abnormalities by a combined VBM and DTI method. This is the first study in which schizophrenia patients have observed no significant abnormality in cerebellar WM volumes, and no significant abnormality in tract-wise WM structural connectivity but revealed decreased FA and increased RD in left MCP and ICP in voxel-wise WM structural connectivity. These findings were consistent with the previous study that evidenced the voxel-based diffusion data analysis is more sensitive than tract-wise analysis in identifying WM abnormalities [31]. Interestingly, our previous meta-analysis study also documented that, compared to HCs, schizophrenia patients exhibited widespread reduced FA but no increased FA[36], and previous study found that the damaged region of WM is mainly revealed in the left side [37]. It should be noted that the cerebellar peduncles (MCP and ICP), as the input fiber of the cerebellum, are the main pathway to communicate with the cerebral cortex and brainstem [38]. Decreased FA and increased RD in left cerebellar peduncles found in schizophrenia patients might be related to the disruption of myelinated axons projecting between the right cerebral cortex and left cerebellar [39]. Besides, neurodevelopmental problems in patients with schizophrenia can lead to abnormalities in brain structure [40]. However, determine whether neurodevelopmental problems are related to FA reduction, this conclusion has not been confirmed. In addition, in VBM, we had not found a significant abnormality in cerebellar WM volume in schizophrenia patients. The study used the diffusion metrics (FA and RD) to determine changes in WM, but not changes in WM using volume (using VBM). Changes in the WM volumes and the diffusion metrics in schizophrenia are not necessarily related [41]. Abnormal FA and RD may reflect changes in myelin and axon membrane integrity, but in general, there are many factors that can regulate FA [42]. In SZ, although FA changes are usually associated with atrophy, they may not have volume changes depending on the method, the region studied and the underlying pathological changes [43]. A meta-analysis comparing WM and DTI changes in schizophrenia showed that DTI studies appeared to be more sensitive to WM abnormalities in schizophrenia [44].

Interestingly, the cognitive assessments were positively correlated with FA and negatively correlated with RD in left cerebellar peduncles in HCs but not in schizophrenia patients. These findings was similar to the previous study that demonstrated the positive correlation between FA in inferior and middle frontal gyrus and cognitive assessments in HCs but not in patients with schizophrenia [45]. Besides, our previous meta-analyses of WM FA in schizophrenia patients showed that the largest abnormal FA was located in bilateral precentral cortex and orbitofrontal cortex, suggesting the cognitive and emotional dysfunction in schizophrenia patients [36]. The impairment of the cerebellar peduncles WM connectivity might induce the pathway aberrant between the cerebellum and the precentral cortex. It can reasonably explain that the reduced FA of abnormality region of the cerebellar peduncles will be strongly related to the abnormal processing speed. Furthermore, the above results also react that reduced FA and a strong relationship with performance on the attention visual task, as well as the interaction of diagnosis by group, shows that this relationship among schizophrenic patients is damaged compared with the control group. Functional imaging studies have always suggested that the dysfunction of the prefrontal cortex, which is the basic structure of higher-order cognitive processing, is an important neural substrate for cognitive dysfunction in schizophrenia (especially during working memory tasks). Our regression linear model showed that cerebellar peduncles predicted attention and working memory behavioral performance in healthy subjects, to elaborate the fact that cerebellar peduncles has an important role in attention and working memory behavioral performance in HCs. However, the above relationship is not significant in schizophrenia patients, showing that the relationship between cerebellar peduncles and cognitive performance may be potential disruption. Furthermore, the decreased FA and increased RD in cerebellar peduncles were associated with disease duration, which evidenced that progressive abnormality of WM integrity in cerebellum in schizophrenia patients.

This study reveals the role of the WM of the brain in the cognitive changes observed in SZ. However, many questions remain unanswered and require further investigation. First, DTI is the most widely used MRI technology available to measure the microstructure of WM. Voxel-based analysis is suitable for analyzing fiber bundles tracing. However, fiber bundles have limitations in cross-cutting, merging, branching, and other regional analyses, and these areas challenge the discussion of examining the cerebellar peduncles [46]. To address these limitations, future DTI studies will need to be conducted using fixel-based analysis [47]. Second, although DTI helps to understand the nature of nerve damage in neurological disorders, the correspondence between DTI indicators and specific underlying pathology remains unclear. Therefore, a decrease in FA and an increase in RD might indicate that white quality changes are caused by demyelination [46]. Subsequent investigations will need to determine whether changes in the white matter of the small brain cause pathological changes in SZ. Finally, we only analyzed the WM structure method and cognitive method of the cerebellar peduncles but did not analyze the functional connectivity and cognitive comparison of WM, so we will calculate both in the future to further describe the structure and function of the cerebellum on the regulation of the cerebral cortical. Future functional imaging studies of patients with brain stem lesions may further understand the role of cerebellar peduncles in cognition.

In summary, we found a statistically significant association between white matter abnormalities and executive dysfunction. Executive dysfunction is one of the most common dysfunctions in the course of schizophrenia [48, 49]. Our regression model showed that the FA of MCP and ICP predicted processing speed, working memory, overall composite and attention vigilance in HCs but not in schizophrenia patients, which might reflect the cognition dysfunction in schizophrenia patients.

## Acknowledgement

Data used in preparation of this article were obtained from the SchizConnect database (http://schizconnect.org) As such, the investigators within SchizConnect contributed to the design and implementation of SchizConnect and/or provided data but did not participate in analysis or writing of this report.

Data collection and sharing for this project was funded by NIMH cooperative agreement 1U01 MH097435

Data was downloaded from the Collaborative Informatics and Neuroimaging Suite Data Exchange tool (COINS; http://coins.mrn.org/dx) and data collection was performed at the Mind Research Network, funded by a Center of Biomedical Research Excellence (COBRE) grant 5P20RR021938/P20GM103472 from the NIH to Dr. Vince Calhoun.

